# Low but significant genetic differentiation underlies biologically meaningful phenotypic divergence in a large Atlantic salmon population

**DOI:** 10.1101/022178

**Authors:** Tutku Aykanat, Susan E. Johnston, Panu Orell, Eero Niemelä, Jaakko Erkinaro, Craig R. Primmer

## Abstract

Despite decades of research assessing the genetic structure of natural populations, the biological meaning of low yet significant genetic divergence often remains unclear due to a lack of associated phenotypic and ecological information. At the same time, structured populations with low genetic divergence and overlapping boundaries can potentially provide excellent models to study adaptation and reproductive isolation in cases where high resolution genetic markers and relevant phenotypic and life history information are available. Here, we combined SNP-based population inference with extensive phenotypic and life history data to identify potential biological mechanisms driving fine scale sub-population differentiation in Atlantic salmon (*Salmo salar*) from the Teno River, a major salmon river in Europe. Two sympatrically occurring sub-populations had low but significant genetic differentiation (*F*_*ST*_ = 0.018) and displayed marked differences in the distribution of life history strategies, including variation in juvenile growth rate, age at maturity and size within age classes. Large, late-maturing individuals were virtually absent from one of the two sub-populations and there were significant differences in juvenile growth rates and size-at-age after oceanic migration between individuals in the respective sub-populations. Our findings suggest that different evolutionary processes affect each sub-population and that hybridization and subsequent selection may maintain low genetic differentiation without hindering adaptive divergence.

## Introduction

Defining populations based on genetic markers has a long history in evolutionary biology (reviewed by Waples & Gaggiotti 2006). The emergence of each new type of molecular marker has seen new discoveries in the extent and scale at which genetic divergence is detected (reviewed by Avise 1994; Wright & Bentzen 1994; Morin *et al.* 2004; Schlotterer 2004). Most recently, studies using single-nucleotide polymorphisms (SNPs) have identified low but statistically significant genetic differentiation in a number of cases where populations were previously thought to be panmictic (O’Reilly *et al.* 2004; Ackerman *et al.* 2011; Zarraonaindia *et al.* 2012; Catchen *et al.* 2013; Garroway *et al.* 2013; Milano *et al.* 2014). Such information is frequently used as the basis for designing management and conservation plans, and in many cases may represent the only information available on population differences. However, the ecological meaning of low but significant genetic differentiation often remains unexplored (Waples & Gaggiotti 2006; Knutsen *et al.* 2011) and relative roles of adaptation, gene flow and the effects of the environment in shaping the genetic structure is not well understood. Likewise, genetically similar populations with dissimilar life histories and morphology may provide insights at the onset of ecological speciation and reproductive isolation (Hendry 2009). Such issues are particularly relevant when considering species or populations of conservation concern and/or harvested species as their interpretation can affect management strategies (Allendorf & Luikart 2007). Integrative approaches, where demographic and phenotypic information are simultaneously assessed alongside genetic analyses, are pivotal for establishing well founded basis for testing ecological-evolutionary hypotheses. However, such breadth of data is often lacking in non-model, wild systems.

Atlantic salmon (*Salmo salar*) is a species of both commercial importance and conservation concern (Verspoor *et al.* 2007). As a result, considerable population genetics research has been conducted on this species, with a variety of molecular markers at various geographic scales (King 2000; King *et al.* 2001; Nilsson *et al.* 2001; Consuegra *et al.* 2002; Verspoor *et al.* 2005; Tonteri *et al.* 2009; Perrier *et al.* 2011; Bourret *et al.* 2013a; Moore *et al.* 2014). Genetic diversity is generally partitioned hierarchically, starting at the continental, followed by basin and then river levels (King *et al.* 2007; Bourret *et al.* 2013a). However, genetic divergence within rivers has also been reported on a number of occasions, where population subdivision at tributary levels are likely to be maintained due to strong homing behaviour (i.e. restricted gene flow) of returning adults and sometimes also local adaptation to different demes (Garant *et al.* 2000; Primmer *et al.* 2006; Dillane *et al.* 2007, 2008; Dionne *et al.* 2008; Olafsson *et al.* 2014).

One of the clearest cases of genetic sub-structuring in wild Atlantic salmon within a river basin has been reported in the Teno River, a large river system in northern Finland and Norway. Microsatellite analyses have revealed surprisingly high levels of genetic divergence across scales of tens of kilometres among tributaries, with average *F*_*ST*_ being around 0.1 (ranging from 0.015 to 0.201; Vähä *et al.* 2007). This divergence was shown to be temporally stable and genetic diversity in the sub-populations was associated with life history variation (Vähä *et al.* 2008). These findings support the notion that sub-populations may be locally adapted. A more recent study using a medium density SNP chip (≈4,300 SNPs) identified several sympatric subpopulation clusters within the river mainstem, with *F*_*ST*_ values at the lower end of those earlier reported (*F*_*ST*_ < 0.0121, Johnston *et al.* 2014). Differences in the distribution of age at maturity (“sea-age”- see below) between sub-population clusters were detected, however, the study focussed on the sea-age phenotype only, and did not include detailed analyses of sub-population structuring within the mainstem and thus the biological significance of the cryptic population structuring remained unclear (Johnston *et al.* 2014).

Sea age at maturity and growth are heritable, complex life-history traits closely linked to fitness in salmonid fishes (Garant *et al.* 2003; Schaffer 2003; Garcia de Leaniz *et al.* 2007; Hutchings 2011; Jonsson & Jonsson 2011). The variation in these traits maintained within and among Atlantic salmon populations are excellent targets for studying evolutionary trade-offs. For example, later maturation at sea is associated with larger size, and therefore higher fecundity in females and higher reproduction success in males, but comes with a cost of higher risk of mortality prior to reproduction (Schaffer 2003). In addition, smaller tributaries with lower water levels are more hospitable to smaller sized, earlier maturing fish, thus providing fitness advantages to younger sea age fish in such tributaries (Garant *et al.* 2003; Niemelä et al. 2006). Likewise, growth, which is inherently linked to several fitness metrics including maturation, survival, and egg size, is likely to be under adaptive constraints associated with intraspecific competition and predator avoidance during juvenile life-history phases (Reid & Peichel 2010; Jonsson & Jonsson 2011), and genetic variation is maintained by context dependent performance in different environments (Gillespie & Turelli 1989; Mackay *et al.* 2009, Reid *et al.* 2012). On the other hand, the underlying genetic and environmental factors shaping reproductive isolation and sea age variation between populations are not well understood. Thus, low genetic differentiation combined with substantial life-history variation within the Teno mainstem populations provides an excellent system for a detailed assessment of whether low but significant genetic differentiation is associated with biologically meaningful phenotypic divergence.

In this study, we utilise the Teno River Atlantic salmon data-set reported in (Johnston *et al.* 2014), and add additional sea age maturity classes and phenotypic data to identify potential biological mechanisms associated with sympatric population divergence in the mainstem of the river. First, we adopted a model-based Bayesian method to refine population structure inference, and subsequently elucidated the spatial distribution of the inferred sub-populations throughout the river. Second, using a wealth of phenotypic and demographic information obtained from fishing records and scale measurements, we provided a detailed account of individual growth rates during different life history stages and demographic properties of each sub-population, and assessed the potential role of natural selection on phenotypic divergence among sub-populations. Our results suggest that despite only subtle genetic divergence, the sub-populations harbour substantial, potentially adaptive, phenotypic divergence including differences in growth rates and size within age classes.

## Materials and Methods

### Study site and sample collection

The Teno River, located in far-north Europe (68–70°N, 25–27°E) runs between Finland and Norway, drains north into the Tana Fjord at the Barents Sea (Figure 1). It supports one of the world’s largest wild Atlantic salmon populations, with up to 50000 individuals being harvested by local fishers and recreational fisheries annually (Johansen *et al.* 2008), accounting for up to 20% of the riverine Atlantic salmon catches in Europe (ICES 2013). A notable feature of the population is the extensive life-history variation observed: age at smoltification (i.e. age of outward migration to sea) varies between two and eight years while the time spent in the marine environment prior to maturation, also called sea-age, varies from one to five years with a proportion of individuals also returning to spawn a second or third time (Niemelä *et al.* 2006). This high diversity of age structure contributes to generally high temporal genetic stability in the system (Vähä *et al.* 2007). Scale samples of returning anadromous adult Atlantic salmon are routinely collected and fish length and weight are recorded by co-operating, trained, fishers within the system. Scales were consistently sampled from below the adipose fin and just above the lateral line (using standard guidelines provided by ICES 2011) and were dried and archived in paper envelopes by Natural Resources Institute Finland (formerly known as the Finnish Game and Fisheries Research Institute). We used the scale sample set reported in Johnston *et al.* (2014) which consisted of fish that return to spawn following one or three to five consecutive winters spent at sea (sea winters, hereafter 1SW (N = 253) and 3SW (N = 283), respectively), and added samples with intermediate maturity time i.e. two sea winter fish (hereafter 2SW, N = 189). A small number of four sea winter (4SW, N= 18) and one five sea winter fish were grouped with the 3SW group (i.e. multi sea winter, MSW); these fish were excluded from growth trait analyses (see below). All fish had been captured along a ~130km stretch of the mainstem Teno River, reaching c. 190km from the sea (Figure 1) between 2001 and 2003. Sampling targeted fish captured during the last 4 weeks of the fishing season in August, which is 2-4 weeks after most individuals have entered the river (Erkinaro *et al.* 2010). As within-river migration to spawning grounds and exploratory movement beyond home spawning areas is limited during the sampling period (Økland *et al.* 2001; Karppinen *et al.* 2004), it is therefore likely that the sampling location is reflective of spawning region in the vast majority of cases. The genetic sex of each fish was determined using the protocol outlined in Yano *et al.* (2013).

**Figure 1:**
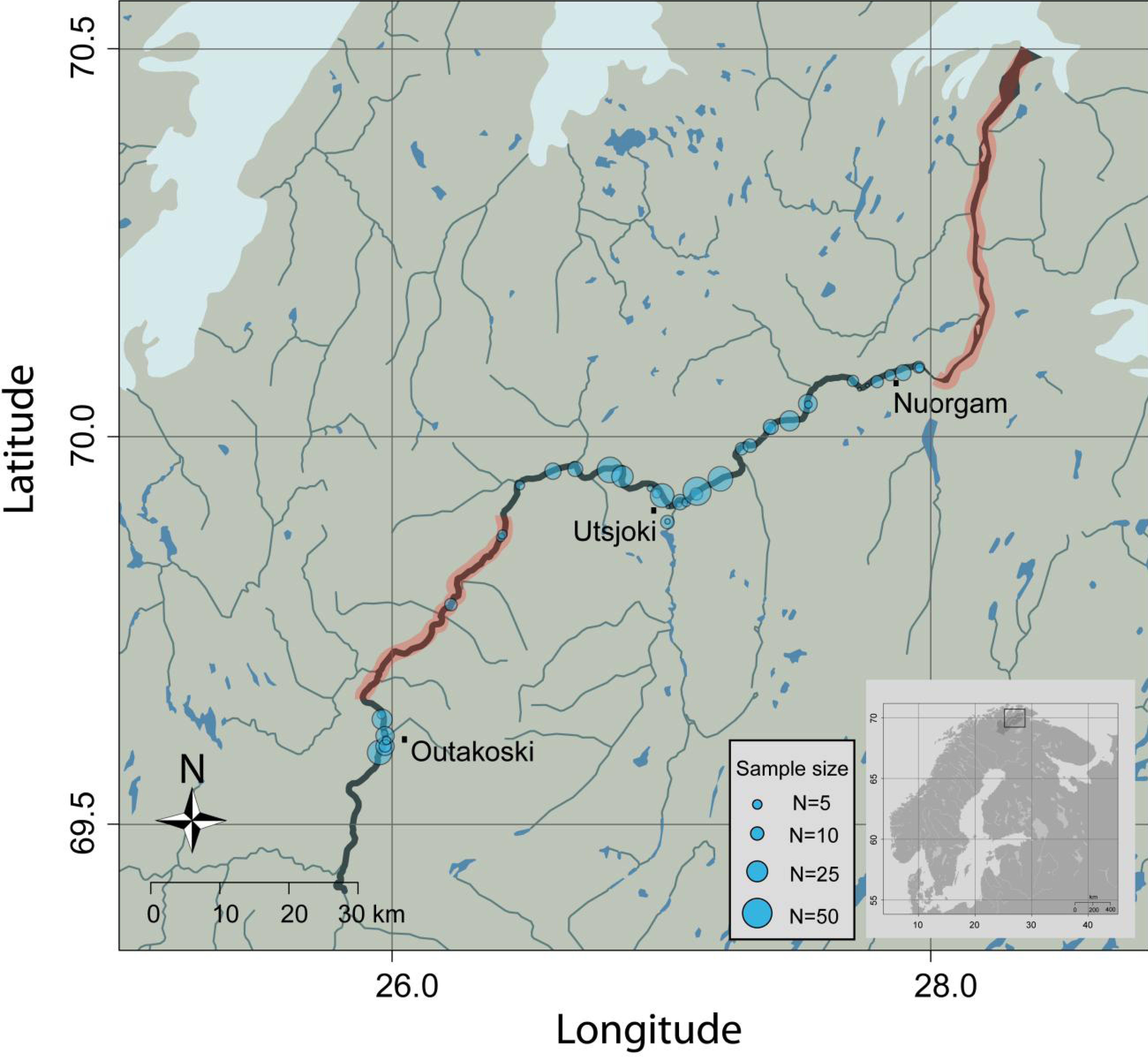
Map of the Teno River and basin with sampling locations along the mainstem (highlighted with a thicker line). Stretches of the mainstem not suitable as spawning grounds or juvenile nurseries are highlighted in red.

### Quantifying morphological and life history traits

We assessed a number of morphological and life history traits extrapolated from scale-derived measurements to determine the biological significance of fine-scale genetic structuring. Scale measures were conducted by trained technicians at the Natural Resources Institute Finland and age and growth rate were determined using the internationally agreed guidelines for Atlantic salmon scale reading (ICES 2011). Seasonal growth variation is reflected in the scale ring patterns, which are used to infer the age of fish (e.g. Friedland & Haas 1996). Likewise, inter-annuli distance (the scale growth between two adjacent annulus rings) is highly correlated to fish growth in the same period (e.g. *r* = 0.96 for juvenile and ocean caught coho salmon (*Oncorhynchus kisutch*), Fisher & Pearcy 1990) and has long been used as proxy for growth rates (e.g. Pierce *et al.* 1996; Erkinaro *et al.* 1997). In the current data, the correlation between total scale growth and adult size was high (Pearson’s *r* = 0.92), and a similarly high correlation is observed between total scale growth in fresh water and freshwater size in a sample set from the same river sytem (Pearson’s *r* = 0.96, Supp. Figure 1). This high correlation between the scale growth and the phenotypes indicates that measurement error should not have a major effect on variance component analysis.

Growth indices were recorded for both the juvenile period (i.e. from the phase in fresh water prior to sea migration) and marine period (feeding phase at sea). In addition to age at smoltification (number of years spent in the fresh water prior to migration to the sea; *FW Age*), several juvenile growth indices were analysed: growth until the end of year one (*Growth*_*FW1,*_ the radius of the scale from the focus to first year annulus), freshwater growth between year one and year two (*Growth*_*FW2,*_ the radius of the scale from the first year annulus to second year annulus), freshwater growth between year two and year three (*Growth*_*FW3,*_ the radius of the scale from the second year annulus to third year annulus), and total freshwater growth (*Growth*_*FWtot*_, scale growth from focus until the end of freshwater growth zone, the point when fish migrates to the sea). In our dataset, all but one individual for which freshwater age data were available spent at least three years in fresh water, therefore *Growth*_*FW1*_, *Growth*_*FW2*_ and *Growth*_*FW3*_ were common metrics for all but one sample. Marine phase indices were: sea age at first maturity (*SW Age*, number of winters spent at sea prior to first migration back to fresh water), first year growth at sea (*Growth*_*SW1*_, the radius of the scale from the end of the freshwater growth to the first year summer annulus). *Growth*_*SW1*_ was the only marine growth parameter that was common to all fish in the data-set. Two terminal traits recorded by the fishers were also included in the analysis; total length at capture (*Length*, i.e. length of the fish from the tip of the snout to the end of the tail) and weight at capture (*Weight*). We also measured body robustness by Fulton’s condition factor at capture (*CF* = 100 × Weight × Length^-3^; Ricker 1975). Phenotypic measurements were available for >90% of samples in all cases except for the yearly freshwater growth parameters (*Growth*_*FW1*_, *Growth*_*FW2,*_ and *Growth*_*FW3*_), which were available for 77% of samples. This was because of the difficultly in confidently assigning annual rings (i.e. annulus) in the freshwater period, which are more prone to scale damage and regeneration of scales.

### DNA extraction, sex determination and genotyping

DNA extraction, sex determination and SNP genotyping for all samples was carried out on individual archived scale samples using the same protocols described in Johnston *et al.* (2014). All 744 samples were genotyped at 5568 SNP loci using a custom-designed Illumina® iSelect SNP-array, the majority of which have been mapped to 29 linkage groups (Lien *et al.* 2011; Bourret *et al.* 2013b). Individual genotypes were scored using the clustering algorithm implemented in the Illumina® GenomeStudio Genotyping Analysis Module v2011.1. Samples with a call rate less than 0.98 were discarded from the analysis. A SNP locus was filtered out if the call rate was less than 0.95, the minor allele frequency (MAF) was less than 0.05 and/or if the heterozygote excess/deficit was significant following false discovery rate adjustment (FDR=0.1), after which 684 individuals remained in the dataset. SNPs in high linkage disequilibrium (LD) were pruned using PLINK’s pruning routine (command *--indep*), using window size=50, sliding window= 5, and variance inflation factor (VIF) = 1.11, the latter corresponding to multiple correlation coefficient of *r*^*2*^=0.1 (Purcell *et al.* 2007). After the pruning step, 2874 SNPs and 684 individuals remained in the dataset. SNPs that were out of Hardy-Weinberg equilibrium were retained, since any population structure may result in HW disequilibrium.

Migrants from distant populations or undetected farmed aquaculture escapees (i.e. among individuals with missing scale growth parameters) were detected from the dataset by calculating pairwise allele sharing between samples using the *ibs* function of the GENABEL package v1.8.0 (Aulchenko *et al.* 2007) implemented in R v 3.1.0 (R Core Development Team 2012). Individuals with average allele sharing distances > 3.09 standard deviations from the median of the distribution (type I error rate probability = 0.001 assuming a normal distribution) were marked as outliers and removed from the analysis. Twenty two (3%) individuals were filtered out at this stage (Supp. figure 2) and a total of 662 individuals remained in the dataset (Supp. table 1).

### Analysis of population structure

Population structure was inferred based on the 2874 SNP markers described above using STRUCTURE Unix version 2.3.3 (Pritchard *et al.* 2000), with 110000 MCMC runs and a burn-in length of 10000, using the correlated allele frequency method (Falush *et al.* 2003) and without defining prior population structure or location. Population structure was inferred by estimating the optimum number of clusters (K) as suggested by Pritchard & Wen (2004) and Evanno *et al.* (2005), in which the smallest K capturing the most structure is concluded as the optimum number of populations explaining the genetic data. K values ranged from one to seven, and each run with a particular K value was replicated 12 times. We then identified each individual’s membership to inferred clusters using a cut off value of q=0.80 (probability of an individual belonging to a group), where q values were averaged over 12 replicated runs. The q=0.80 threshold is conservative for assigning individuals to populations (see Vähä & Primmer 2006), and also allows the distinction of some hybrid classes from pure-breds (e.g. backcross hybrids are expected to have a q-value around 0.75; see below). Individuals not assigned to any population cluster (q-value < 0.80) were defined as “admixed”.

Following population inference, Weir’s and Cockerham’s pairwise *F*_*ST*_ (Weir & Cockerham 1984) and within sub-population genetic diversity indices (i.e. observed and expected heterozygosity) were estimated within and among the inferred sub-populations using the HIERFSTAT package v0.04-10 (Goudet 2005) in R v3.1.0. Diversity indices of inferred sub-populations were compared with Kruskal-Wallis test.

### Demographic and phenotypic properties of sub-populations

To evaluate genetic isolation by distance in the data-set, associations between individual-level genetic distances (i.e. allele sharing) and geographic distances (i.e. approximate river position) were assessed using a Mantel test, and significance was evaluated by permuting the data 10,000 times using the VEGAN package v2.0-10 in R v 3.1.0 (Oksanen *et al.* 2013). In addition to isolation by distance, we also tested for a possible isolation by region signature along the lower and the upper section of the mainstem, which are separated by a 40 km stretch of sandy river habitat that is generally unsuitable for salmon reproduction and nursery (Niemelä *et al.* 1999, Figure 1). Because of this, we also included a test of genetic isolation by region where genetic similarity of fish from the lower (< 140km) and the upper (> 180 km) stretches of the river were compared. A small number of fish sampled within this sandy region (3% of the final dataset) were excluded from this Mantel test. We constructed the distance matrix as follows: any two fish that were sampled in the same region were scored as “0” in the distance matrix (i.e. no distance between them), whereas fish that were not sampled in the same region were scored “1”. Finally, we quantified the relative contribution of distance (km) vs sub-region (upper vs lower) effect in explaining the pairwise genetic distance between individuals. The two matrices (distance matrix vs sub-region matrix) are inherently confounded, thus we used a partial Mantel test to identify the relative contribution of each one, in which the correlation between the genetic distance matrix and either of the spatial matrices are conditioned on the other spatial matrices (using *mantel.partial* function in the VEGAN package v2.0-10). Significance was assessed at alpha value of 0.00625, after Bonferroni correction for multiple testing.

The among sub-population variation in continuous growth traits was evaluated using a linear mixed effect model, where parameters were estimated with maximum likelihood using the LME4 package v1.1-7 in R v 3.1.0 (Bates 2010). The model included the sub-population of origin (as inferred by structure analysis at q = 0.8), *SW age*, *FW age*, and the genetically assigned sex as fixed effects, and year of sampling as a random effect. These covariates were chosen because they are either inherently or likely to be associated with the traits of interest. For example, *SW age* and sex are both strong predictors of sea growth, while *FW age* is a good predictor of freshwater growth and total size in the fresh water. The model was parametrically bootstrapped 10000 times using the *bootMer* function in LME4, from which the sampling median and 95% confidence interval of the parameters were calculated. Finally, the null hypothesis, that the parameter has no effect on the response variable, was evaluated at two alpha values, 0.05 and 0.001, which denote the proportion of (bootstrapped) parameter estimates with an opposite sign to the null. All phenotypic measurements other than *CF* were log scaled to achieve normality. In addition to the continuous traits, the two categorical traits *FW age* and *SW age* were tested for association with sub-population of origin, using a generalized linear model (Poisson error function and log link), where *SW age* was modelled as number of years that maturation was delayed beyond *SW age* = 1, otherwise with the same procedure as above. We then extended the phenotype analysis to assess a potential isolation barrier between the upper and lower sections of the river that are separated by a sandy stretch of river that is mostly unsuitable for spawning and juvenile rearing. Therefore, we reformulated the above linear mixed effect by replacing the “sub-population” term with “sub-population and region” effect, where each sub-population and region combination was accounted as a categorical fixed effect in the model. Similar to the previous model, the parameter confidence intervals were estimated by parametric bootstrapping with 10000 permutations.

### Genome wide association with phenotypes

Genome wide association studies (GWAS) were performed on all 10 phenotypic traits outlined above. Eight continuous traits were modelled using general linear models with a Gaussian error structure, fitting SNP genotypes and all covariates significantly associated with the response variable as fixed effects; two traits (FW Age and Sea Age) were modelled using Poisson function as the link, where *SW age* was modeled as number of years maturation was delayed beyond *SW age* = 1. A GWAS of 1SW vs 3-4SW individuals was conducted earlier (Johnston *et al.* 2014), however here a larger data-set including 2SW individuals and additional phenotypic traits was investigated. Population stratification was accounted for either by including the significant principal components to the model as fixed effects, or using genomic control whereby the test statistic was divided by the genomic inflation factor (i.e. λ, Price *et al.* 2010). Principal components were added sequentially until the inflation factor (lambda) was less than 1.1. The significance threshold for genome-wide association after multiple testing at α= 0.05 was calculated using the Bonferroni method.

### Adaptive divergence among populations

We evaluated the role of adaptive divergent selection among populations using a *P*_ST_–*F*_ST_ comparison (Brommer 2011). This is an extension of the *Q*_ST_–*F*_ST_ framework, in which the proportion of additive genetic contribution to population divergence is estimated within a range of values to infer the robustness of the selection signal. This was determined using the following equation:

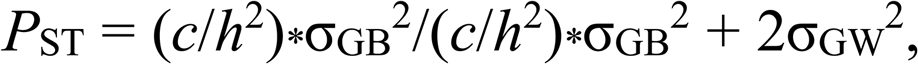

where σGB^2^ and σ GW^2^ are the variances between and within each population, respectively (i.e. residuals of the model); *h*^2^ is heritability; and *c* is the proportion of the total variance that is presumed to be due to additive genetic effects across populations (Leinonen *et al.* 2006; Brommer 2011). We estimated the among population variation using a mixed model approach, where significant covariates (as evaluated in the linear model above) were included as fixed terms and population provenance as a random term using a restricted maximum likelihood approach (REML) as implemented in the LME4 package v1.1-7 (Bates and Maechler 2009) in R 3.0.2 (R Core Team). *FW Age* and *SW Age* were fitted using a generalized model with a Poisson link, where *SW age* was modeled as number of years maturation was delayed beyond *SW age* = 1. In this analysis, we included only individuals that were confidently assigned to a population (q>0.8). Finally, models were bootstrapped 10000 times using the *bootMer* function in LME4 (with *use.u=T* option), from which the confidence interval of the parameters were calculated. We calculated *F*_ST_ distribution by performing an *F*_ST_-outlier analysis in ARLEQUIN 3.5 (Excoffier *et al.* 2005, Beaumont & Nichols 1996). The highest non-significant *F*_ST_ value at *a* = 0.05 was taken as the upper threshold for the neutral expectations. In natural populations, the empirical values of *c* and *h*^2^ are often unknown; therefore, we tested the robustness of *P*_ST_–*F*_ST_ comparisons within a specified range of *c*/*h*^2^ ratios (0 to 2) as recommended by (Brommer 2011).

### Estimating admixture between the inferred sub-populations

In order to gain further insight into the patterns of gene flow among sub-populations (e.g. Taylor 2003), we estimated the composition of different hybrid classes within the admixed individuals. To do this, we used the q-value of an individual as a proxy for its hybrid index (Vähä & Primmer 2006). First, we assessed the expected q-value distribution of different hybrid classes by simulating individuals using the empirical frequency distribution of inferred sub-populations. We simulated three different hybrid classes, assuming no linkage: 1) F_1_ hybrids; 2) F_2_ hybrids (i.e. F_1_ × F_1_); and 3) Backcross hybrids (F_1_ × pure-bred sub-population 1 or 2). A baseline of pure type individuals (N = 400 for each population) was generated by sampling the observed allele frequency distributions (using genotypes inferred in the population structure analysis), and the population of origin for this group were marked *a priori* in the STRUCTURE analysis (using POPFLAG = 1). Next, 200 individuals from each hybrid class were simulated and q-value distributions were retrieved using STRUCTURE software using the same parameters as above. The q-value distributions of simulated hybrid classes were visually compared to the distribution of empirical q-values in order to infer the possible hybrid structure within the empirical data.

## Results

### Analysis of population structure

The STRUCTURE analysis showed a rapid increase in the log likelihood value from K=1 to K=2, followed by a plateau (Figure 2a), suggesting K=2 as the optimal number of sub-populations identified within the genetic data. This conclusion was also supported by the Δ K method of (Evanno *et al.* 2005), where Δ K was highest at K=2 (Supp. Figure 3). Using a conservative q-value threshold of 0.80 (see Materials and Methods), 52% (N = 347) and 26% (N = 171) of individuals were assigned to the two main clusters, whereas 22% (*N* = 144) were assigned as admixed (Figure 2b, Supp. figure 4). Therefore, we refer to these two distinct sub-populations as “Sub-population 1”, and “Sub-population 2” hereafter, while the remaining samples are referred to as “admixed”. Individuals assigned to clusters in the STRUCTURE analysis also grouped together in the principle component analysis (PCA), where the first two principle component (PC) explained 6.7 and 5.6% of the genetic variation respectively (Figure 2c).

**Figure 2:**
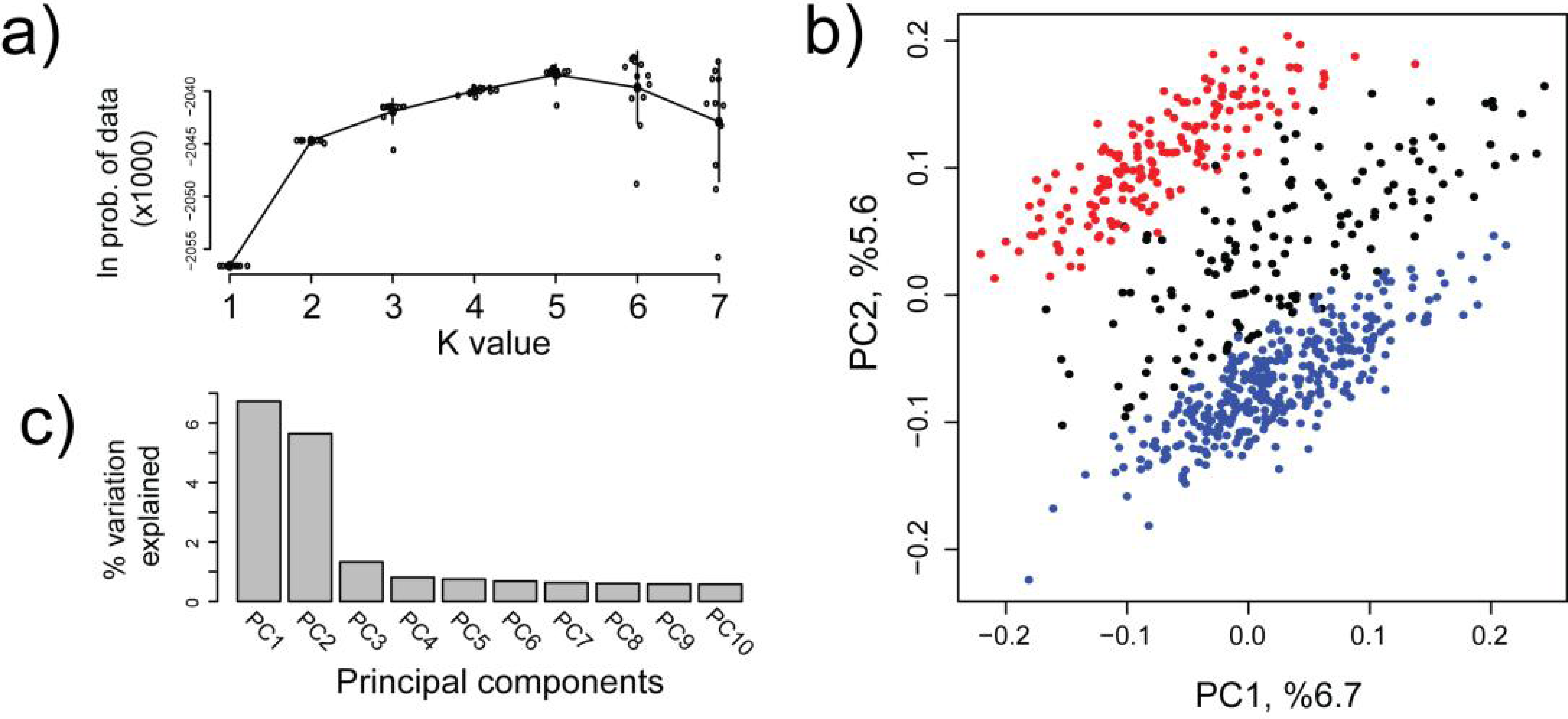
STRUCTURE and principal component analyses of Atlantic salmon sampled from the Teno River mainstem. a) The estimated *ln* probability of data given the K value. Error bars are standard deviations of 12 replicate runs. The results for each of the 12 replicate runs are given with smaller circles; b) Plot of the first two major PC axes, where colors show sub-populations inferred by the STRUCTURE analysis at the optimum *K* value of two. Blue, red and, black colors show Sub-population 1, Sub-population 2, and admixed individuals, respectively. (c) Percent variation explained by the first 10 PC axes.

**Figure 3:**
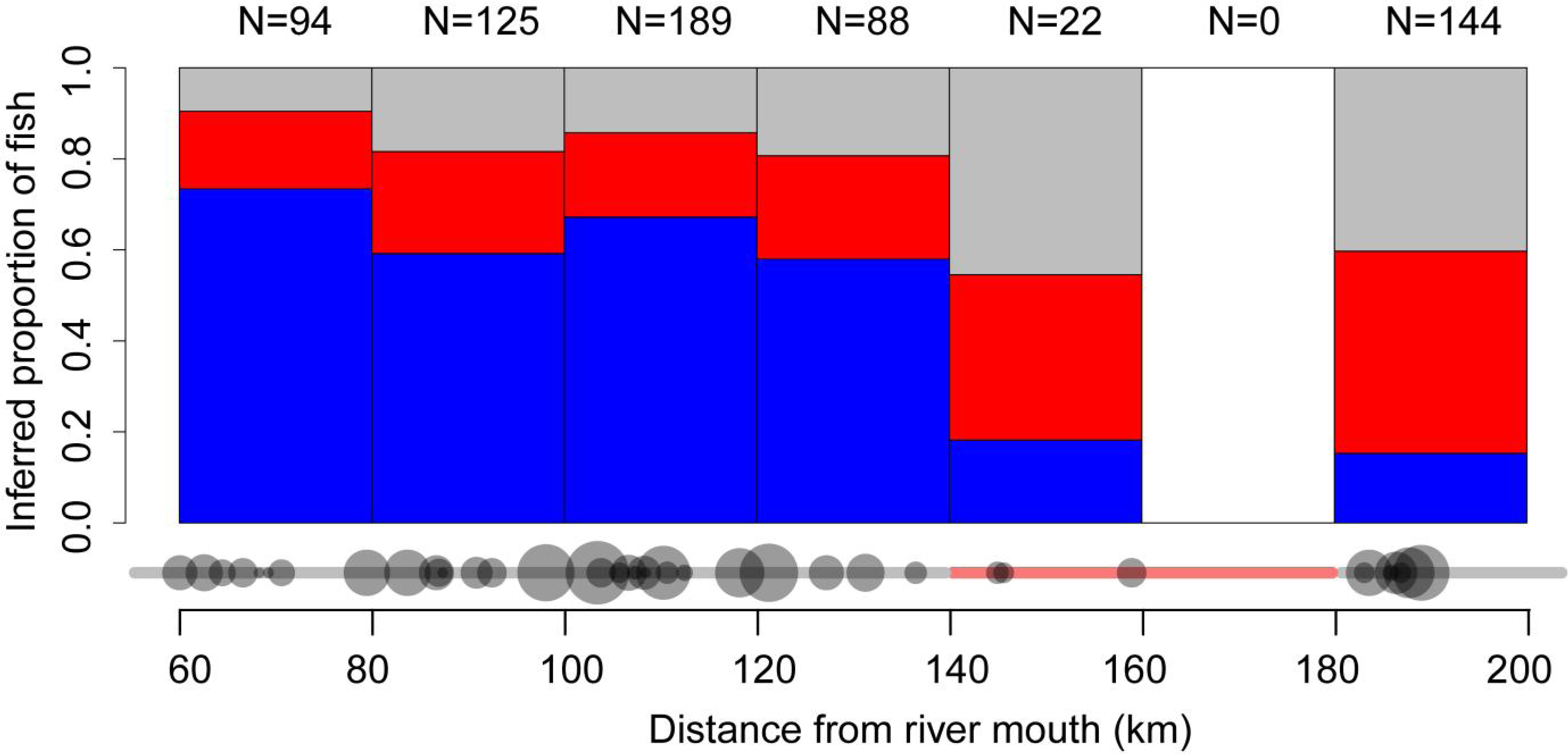
Proportions of the inferred Atlantic salmon sub-populations over the sampling range along the Teno River mainstem. Blue, red and, grey colors indicate Sub-population 1, Sub-population 2, and the admixed group, respectively. Proportional sample sizes for specific locations along the mainstem are indicated by circle diameter and total sample sizes within 20km intervals are listed above the bars. The sandy stretch of river that is mostly unsuitable for spawning and juvenile rearing is indicated with red on the lower horizontal line.

**Figure 4:**
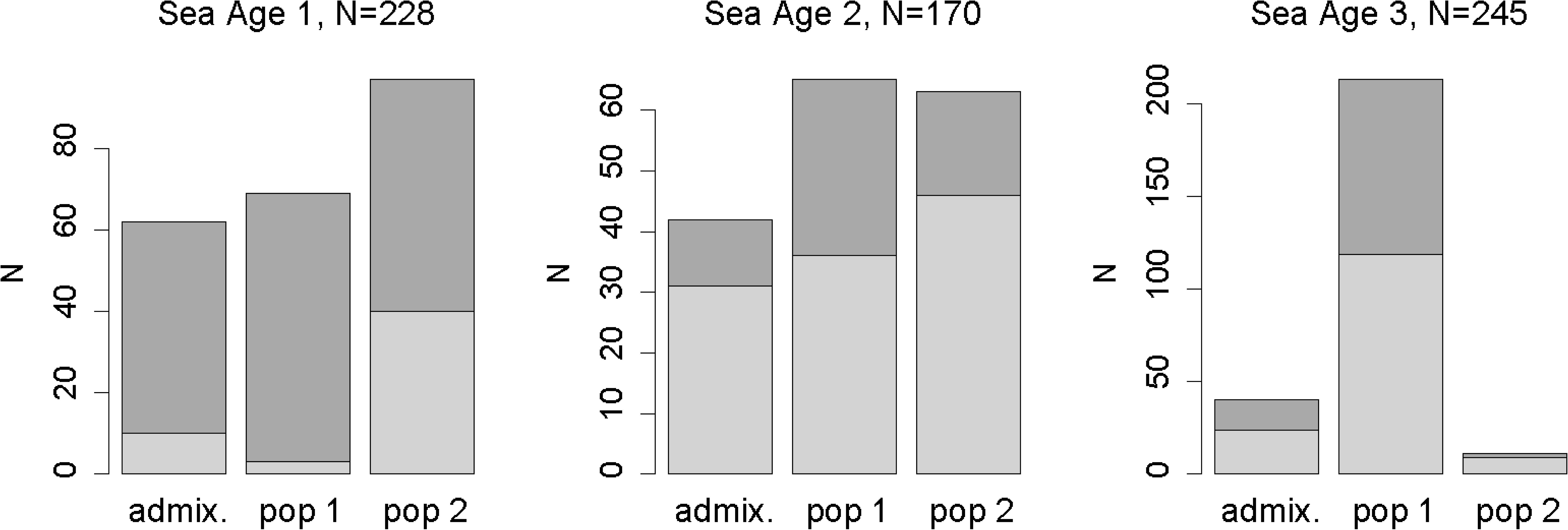
Sex distribution (males in dark-grey, females in light-grey) among sub-populations and sea age classes of Atlantic salmon in the Teno River mainstem.

Expected and observed heterozygosity was marginally but significantly larger in Sub-population 1 compared to Sub-population 2 (Kruskal-Wallis test, Table 1). Genetic differentiation between the two sub-populations was *F*_*ST*_ = 0.018 (95% CI= 0.017 -0.019, Weir and Cockerham’s *F*_*ST*_).

**Table 1:** Diversity indices (mean, sd) of the mainstem Teno River Atlantic salmon clusters inferred by STRUCTURE (*N*_*SNP*_= 2684). *H*_*O*_ =observed heterozygosity, *H*_*O*_ = genetic diversity, *F*_*IS*_.= inbreeding coefficient

| | $H_O$ | $H_S$ | $F_{IS}$ |
| --- | --- | --- | --- |
| Sub-population 1 | 0.3588 (0.1205) | 0.3604 (0.1195) | 0.0040 (0.0538) |
| Sub-population 2 | 0.3518 (0.1300) | 0.3546 (0.1281) | 0.0076 (0.0779) |
| Admixed | 0.3559 (0.1214) | 0.3638 (0.1195) | 0.0214 (0.0861) |

### Demographic and phenotypic properties within and among the inferred sub-populations

Fish from distinct sub-populations were not distributed evenly, nor grouped completely separately along the sampled stretch of the mainstem. A higher proportion of Sub-population 1 fish was present in the lower Teno, and Sub-population 2 fish were more common in the upper Teno (Figure 3). There was a marked change in the proportions of sub-populations around the river stretch that is unsuitable for spawning after c. 130 km (Figure 3). There were no significant differences in sampling time between populations, sea age or their interaction, suggesting both populations, and the different sea age groups within them, are likely to have similar spawning periods (Supp. table 2).

Individual-level isolation by distance (IBD) within Sub-population 1 revealed a marginal but non-significant signal after the multiple test correction (Mantel’s *r*= 0.063, *p* = 0.007; Table 2). Sub-population 2 showed slightly weaker IBD patterns (Mantel’s *r* = 0.032, *p* = 0.020; Table 2). The isolation by region analysis testing for genetic isolation between upper and lower Teno mainstem samples was significant for Sub-population 1 (Mantel’s *r*= 0.093, *p* = 0.002), but not for Sub-population 2 (Mantel’s *r* = 0.036, *p* = 0.018). Partial Mantel tests, by which confounded effects of linear distance and region on genetic distance were partitioned, suggested that the genetic divergence in Sub-population 1 was driven primarily by restricted gene flow between regions (Mantel’s *r*= 0.075, *p* = 0.001, Table 2), but this was not the case in Sub-population 2 (Mantel’s *r*= 0.006, *p* = 0.455, Table 2), suggesting lack of divergence between upper and lower Teno fish from Sub-population 2.

**Table 2:** Isolation by distance analyses in the two Teno River Atlantic salmon sub-populations.

| Sub-population | $N^a$ | distance matrix | Partial Mantel correction matrix<sup>2</sup> | Mantel's $r$ | $p$ value <sup>b</sup> |
| --- | --- | --- | --- | --- | --- |
| Sub-population 1 | 347 | distance |  | 0.063 | 0.007 |
|  | 347 | sub-region |  | 0.093 | **0.002** |
|  | 347 | distance | sub-region | -0.021 | 0.892 |
|  | 347 | sub-region | distance | 0.075 | **0.001** |
| Sub-population 2 | 171 | distance |  | 0.032 | 0.020 |
|  | 171 | sub-region |  | 0.036 | 0.018 |
|  | 171 | distance | sub-region | 0.011 | 0.356 |
|  | 171 | sub-region | distance | 0.006 | 0.455 |

<sup>a</sup> Number of individuals in the analysis. <sup>b</sup> Bold letters indicate significant $\alpha$ values after multiple test correction; $\alpha= 0.00625$

There were striking differences in the proportion of sea age classes assigned to each sub-population and the sex-ratios within each population (Figure 4). Most 3SW fish were assigned to Sub-population 1 (88% of 264 3SW fish) while only 11 (4%) were assigned to Sub-population 2. Almost all 1SW fish assigned to Sub-population 1 were male (78 of 82 fish, 95%). In contrast, there were no apparent differences in the distribution of 2SW fish between sub-populations (Figure 4). No difference in the freshwater age distribution was observed between sub-populations, nor was there any association between sea age and freshwater age (Table 3).

**Table 3:**
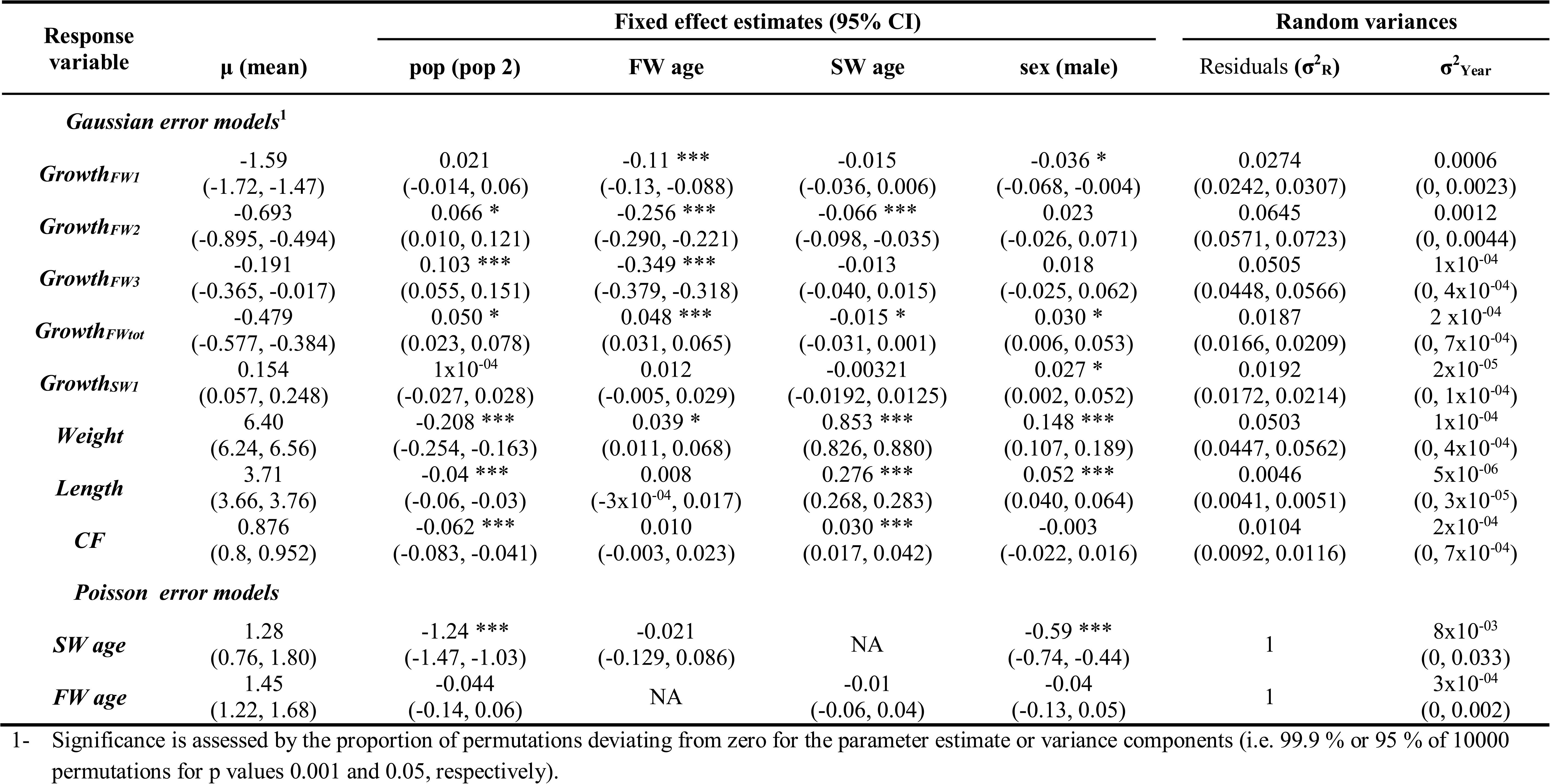
Estimated fixed effects and random variance components in the mixed model analysis of phenotypic variation within and between the inferred populations of Atlantic salmon in the mainstem Teno River. The 95% confidence intervals, estimated by parametric bootstrapping, are given in parentheses. Asterisks denote effect sizes significantly different from zero^1^ (*** = 0.001, *= 0.05).All continuous traits other than condition factor are log scaled

Continuous growth traits were also significantly different between sub-populations. Out of nine growth/size traits measured, six showed significant differences between sub-populations (Table 3, Figure 5). In general, freshwater growth rate was faster for Sub-population 2, however, following the marine period, this was reversed and at the time of sampling, fish from Sub-population 1 were significantly larger in length and weight, and had higher condition factors than Sub-population 2 individuals (Figure 5, Table 3). For example, the average weight differences between individuals of the same sea age classes from Sub-populations 1 and 2 were 0.21 kg (11%) and 3 kg (34%) for 1SW and 3SW fish, respectively (see Table 3 for parameters and log scale CIs). Sex was a significant determinant for growth at sea traits, such that males grew more in the first year at sea (*Growth*_*SW1*_) and were longer and heavier at return (Table 3). Males had also grown more by the end of the freshwater period (*Growth*_*FWtot*_, Table 3). Finally, higher sea age at maturity (*SW Age*) was significantly associated with slower freshwater growth (*Growth*_*FW2*_ and *Growth*_*FWtot*_; Table 3).

**Figure 5:**
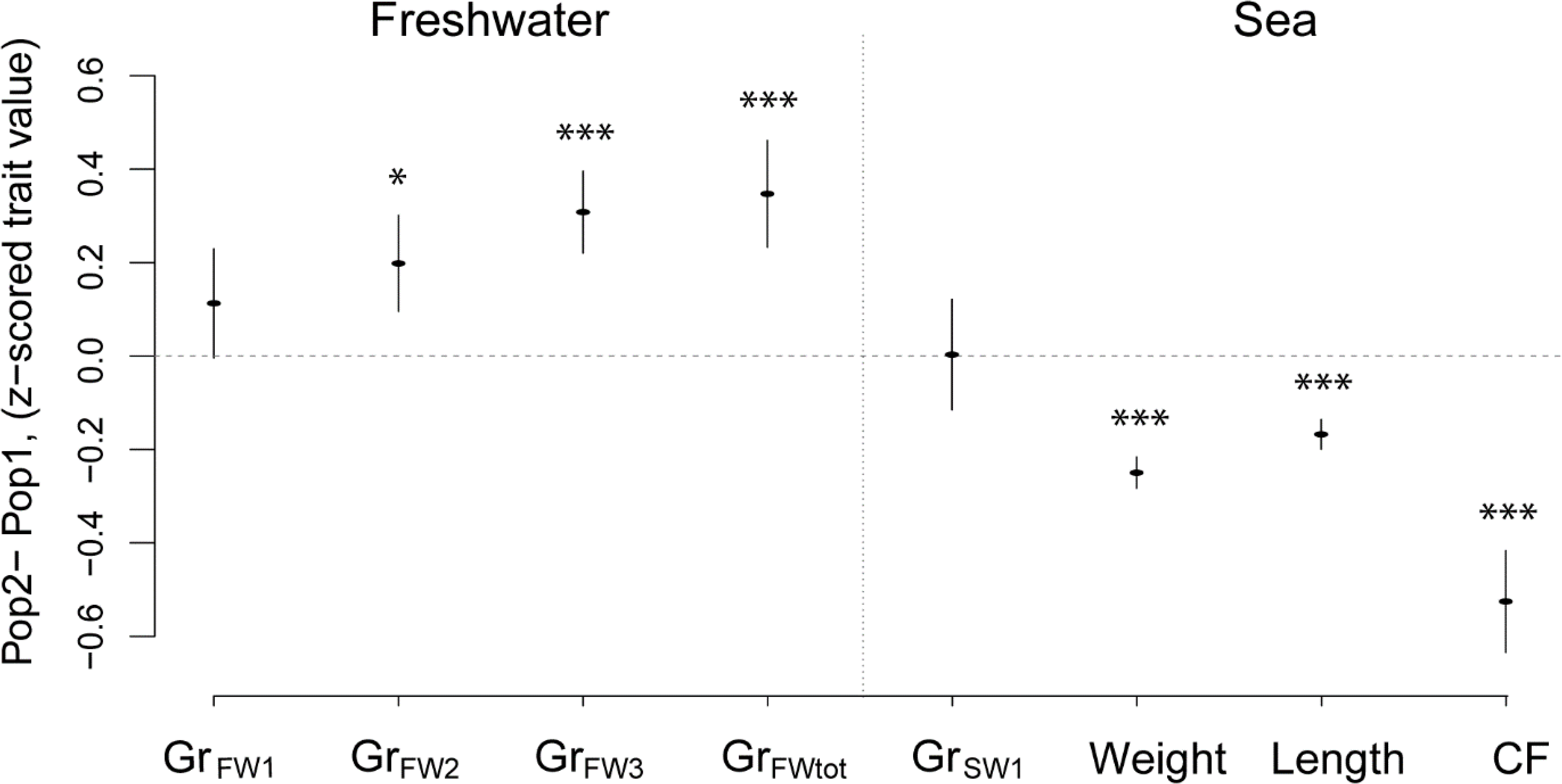
Population specific differences between phenotypic trait values of Teno River Atlantic salmon sub-populations. Bars shows standard deviations of the differences inferred from 10000 permutations. Asterisks denote significant differences between populations (*** = 0.001, *= 0.05). Here, only population specific effects are accounted for after being inferred by the linear model (See Table 2 for details).

When sampling location was taken into account, we observed significant differences in the freshwater growth trajectories between sub-population 1 individuals from the upper and lower mainstem regions with higher growth in the upper region (i.e. *Growth*_*FW2*_ and *Growth*_*FWtot*_ in Supp. figure 5). However, this fast growth appears to slow down in the first year at sea (i.e. *Growth*_*SW1*_) and both upper Teno and lower Teno Sub-population 1 attain similar size at return (Supp. figure 5). Unlike Sub-population 1, Sub-population 2 fish sampled in the upper and lower Teno exhibited similar growth both in the fresh water and in the sea (Supp. figure 5).

### Genome wide association studies

None of the 2874 SNP loci showed a genome-wide significant association with any trait, after correction for population stratification using the principal component method (Price *et al.* 2006). A number of SNPs were significant at a non-conservative alpha value of 0.01, but allelic substitution effects of these SNPs did not explain phenotypic variation within sub-populations, more than by chance alone (Supp. figure 6), further indicating that these loci are likely false positives. The only exception was the condition factor, where 3.9% and 6.6% of phenotypic variation were explained by the top 28 significant SNPs in each sub-populations respectively (*p*<0.01), suggesting a small polygenic effect on condition factor can be explained by these SNPs (Supp. figure 7. See figure legend for details). When using the genomic control method alone to account for population stratification, a significant association between several genome regions and *SW Age* was observed; this is consistent with the significant genome regions identified Johnston et al. (2014) when comparing 1SW and 3SW fish using the genomic control method, but not with correction using principle components or when modelling identity-by-state between individuals. However, as acknowledged in the previous study, effective population sizes within the Teno mainstem are high, whilst genome-wide levels of linkage disequilibrium are low. Therefore, we cannot rule out that absence of associations are due to low heritability and/or a polygenic basis of these traits, or if marker density and sample size are insufficient to capture variation at markers in strong linkage disequilibrium with causal variants (see discussion in Johnston et al. 2014).

### Adaptive divergence among populations

The phenotypic differences between sub-populations in terminal traits including *Length*, *CF*, and *Sea Age*, were consistent with selection contributing to the divergence, whereby *P*_ST_ estimates and 95% CI of these traits were larger than neutral range, which was also robust across a wide range of *c*/*h*^*2*^ values (Figure 6). *P*_ST_ estimates for these traits remained above the neutral range at *c*/*h* values as low as 0.5, suggesting that trait variances may be subjected to divergent selection even when the proportion of additive genetic affect among populations is half of the within population value (see Brommer 2011). Population variance between freshwater traits was not significantly different from neutral expectations, although median *P*_ST_ estimates for juvenile growth during later years in the river (i.e. *Growth*_*FW3*_, and *Growth*_*FWTot*_) was larger than the neutral range at higher *c*/*h*^*2*^ values, weakly suggesting divergent selection may potentially influence these traits.

**Figure 6.**
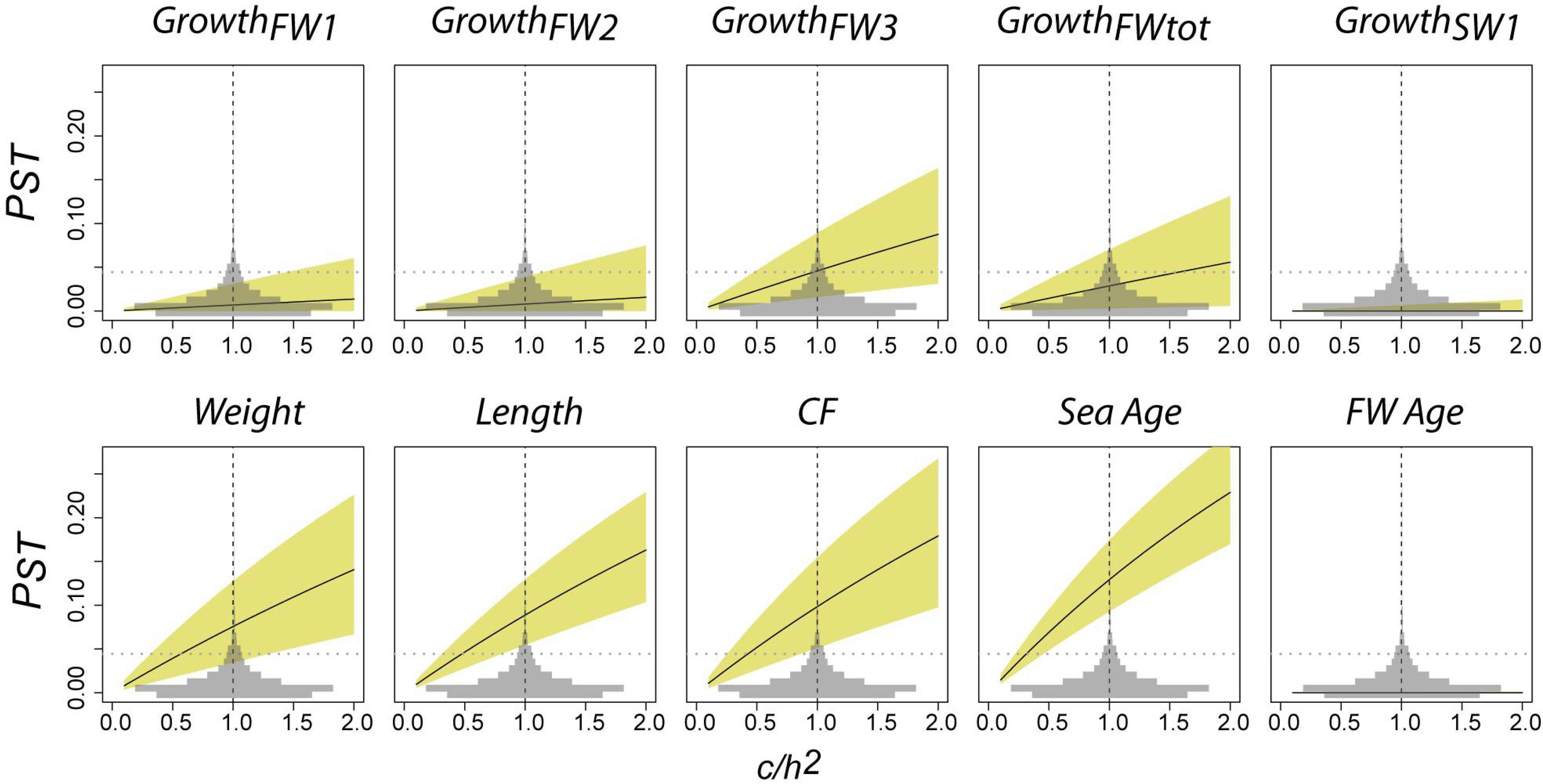
The relationship between *P*_ST_ and *F*_ST_ between the two Teno mainstem Atlantic salmon populations under different *c*/*h*^2^ ratio scenarios for the 10 phenotypic traits assessed in this study. The SNP *F*_ST_ distribution is plotted in light grey and the upper neutral *F*_ST_ estimate is indicated with a grey horizontal line within each plot. The vertical dashed line in each panel shows the *c*/*h*^*2*^ value at 1, where the relative contribution of additive genetic effects to population variation (*c*) is equal to (*h*^*2*^). The median *P*_ST_ estimate is shown with a solid black line, and the coloured area indicates the 95 % CI of the *P*_ST_ estimate.

### Population admixture between the inferred sub-populations

A substantial proportion of sampled fish (21.8%; Figure 2b and Figure 3) had intermediate q-values, suggesting that admixture in the system was common. The empirical q-value distribution of admixed fish was skewed towards Sub-population 1 which suggests genomes from admixed individuals contain a higher proportion of alleles from Sub-population 1 (Supp. figure 8). The relatively “flat” distribution of q-values suggests that the admixed individuals also include higher order hybrids (Supp. figure 8). On the other hand, the admixed group had high *F*_*IS*_, which cannot be explained by inbreeding (i.e. overall high *Ho* of the group, see Table 1). However, a heterogeneous origin of populations within a group would elevate the *F*_*IS*_ signal, suggesting some fish in the admixed group may have origins other than the two sub-populations in the study, perhaps other sub-populations from other tributaries in the Teno River system.

## Discussion

We combined SNP-based sub-population inference with extensive phenotypic and life history data to obtain a detailed account of fine-scale population differentiation in Atlantic salmon from the mainstem of the Teno River, a major salmon river in Europe. Our results suggest that despite only subtle genetic divergence (*F*_*ST*_ = 0.018), the two sub-populations do indeed harbour substantial phenotypic divergence, including differences in age structure, growth rates and size within age classes. Although both sub-populations inhabited overlapping sections of the river, Sub-population 2 appeared to have a broader range extending towards the upper Teno mainstem. This suggests that different evolutionary processes may maintain divergence between these two genetically similar, overlapping sub-populations. Furthermore, strong signatures of adaptive divergence at sea, coupled with seemingly similar spawning timing and location leave open the possibility of a link between reproductive isolation and divergence at sea. In this discussion, we consider the potential processes that may be driving this population structuring, as well as the broader significance of the findings from both evolutionary and conservation management perspectives.

### Partial reproductive isolation in sympatry: possible mechanisms

Detailed spatial analyses indicated that members of each sub-population were distributed throughout the mainstem of the river, suggesting that the two sub-populations occur in sympatry. Reproductive isolation in sympatry between populations of the same species or closely related species provide good study systems for understanding the evolution of reproductive isolation, and hence ecological speciation (e.g. Huber *et al.* 2007; Nosil & Sandoval 2008; Stelkens *et al.* 2010; Arnegard *et al.* 2014. See also, Hendry 2009). Due to their current habitats being in previously glaciated regions, salmonid fishes have frequently been the focus of studies investigating the mechanisms involved in the early stages of ecological speciation. However, in the vast majority of these cases, reproductive isolation between populations is mediated by extensive dichotomy in life history variation: examples include anadromous vs resident strategies in Atlantic salmon (Verspoor & Cole 1989; Vuorinen & Berg 1989) and steelhead/rainbow trout, *O. mykiss* (Docker & Heath 2003; Narum *et al.* 2004; Pearse *et al.* 2009; Hecht *et al.* 2013); run timing variation in pink salmon, *O. gorbuscha* (Gharrett *et al.* 2013); and freshwater (kokanee) and marine (sockeye) migrating populations of *O. nerka*, (Taylor 1999). Likewise, species pairs with diverged ecotypes, which may have overlapping breeding ranges, show discontinuous adaptive variation and strong genetic differentiation as a result of established post- and pre-zygotic reproductive isolation (Gislason *et al.* 1999; Taylor 1999; Saint-Laurent *et al.* 2003; Østbye *et al.* 2005; Landry *et al*. 2007; Hendry 2009; Power *et al.* 2009; Kapralova *et al.* 2011; May-McNally *et al*. 2015).

In comparison, the results reported here provide a novel case of phenotypic divergence between populations with very subtle genetic divergence, where gene flow between populations is restricted despite an overlapping breeding range, similar basic life histories (e.g. both sub-populations are anadromous) and similar spawning periods. The potential mechanisms maintaining the population structure are therefore less clear than in some earlier cases. In our study, both sub-populations exhibited skewed age structure between sexes, where males mature earlier, spending fewer years feeding at sea. This is consistent with previous work (e.g. Fleming 1998; Niemelä et al. 2006), and is likely a result of the tighter positive correlation between reproductive output and increasing size, and hence age, in females, compared to males (Fleming 1996; Fleming 1998). On the other hand the difference in sea age structure between the genetically similar sub-populations is curious. Below, we consider potential pre- and post-zygotic isolation mechanisms that could potentially lead to the observed genetic and phenotypic divergence.

A potential pre-zygotic reproductive isolation mechanism is micro-geographic separation of spawning areas throughout the mainstem Teno River. It is known that breeding site preference in Atlantic salmon is partly driven by gravel size (Louhi *et al.* 2008), whereby areas with faster flowing water and larger gravel size are only accessible to larger females (Fleming & Einum 2010). Given that Sub-population 1 is essentially devoid of small, 1SW females, whereas Sub-population 2 almost completely lacks large 3SW females, size-assortative breeding site selection could provide the means for at least partial reproductive isolation on a micro-geographic scale. On the other hand, this argument does not explain the genetic divergence satisfactorily, since 2SW females are relatively common in both sub-populations (Figure 4), and size-assortative breeding sites of females may not restrict gene flow via males. Moreover, gravel size is not known to be different in the upper and lower section of the mainstem (J. Erkinaro, *unpubl. data*).

Inference of possible post-zygotic reproductive mechanisms assumes that there is a fitness disadvantage for hybrid individuals (Turelli *et al.* 2001; Servedio & Noor 2003), which in turn requires the assumption that the two sub-populations are locally adapted. Although the relatively flat distribution of admixed q-values observed here suggests that the admixed fish can survive and reproduce for more than few generations, there is some circumstantial evidence that could provide a basis for post-zygotic isolation if the sub-populations are indeed locally adapted. Firstly, size at return from the marine migration is significantly different between sub-populations, and consistent with adaptive divergence. For example, 3SW female fish from Sub-population 1 are ~9.9 kg in weight (N=108) compared to ~7.6 kg for the few 3SW fish from Sub-population 2 (N=9, see also Table 3 for parameters and log scale CI). Likewise, 2SW and 1SW fish from Sub-population 1 (N = 65 and 69, respectively) are about 2.0 kg and 0.25 kg heavier, respectively, than comparable fish from Sub-population 2 (N = 63 and 97), after adjusting for sex. In addition to size, condition factor is also significantly different between sub-populations, with fish from Sub-population 1 having a higher condition factor on return from the sea (Figure 5). This dramatic difference in size and condition of fish following the marine feeding phase could be explained either by the sub-populations exploiting different marine feeding grounds, or by differences in their efficiency to exploit the same feeding grounds. Very little is known about the marine feeding phases of most salmon populations (Haugland *et al.* 2006; Chaput 2012; MacKenzie *et al.* 2012), and thus this issue requires further research. Nevertheless, the pronounced size difference in returning adults provide a plausible post-zygotic isolation mechanism if the marine feeding strategy/behaviour of hybrids was sub-optimal, and therefore hybrids had lower survival compared to the pure-breds of either sub-population. The high *P*_ST_ values in these traits is also consistent with divergent selection in the marine environment (Figure 6) thus further supporting the significance of the marine habitat for population structure.

### Faster freshwater growth – earlier sea age at maturity

Our results suggest Sub-population 1 was mostly confined to the lower Teno mainstem, while Sub-population 2, which seemingly performed poorer at sea, was inhabiting the entire sampling range of the mainstem. Intriguingly, even in the lower mainstem where individuals of the two sub-populations occur sympatrically, individuals of Sub-population 2 had higher growth in the fresh water, suggesting the growth differences are not due to spatial geographical variation (Supp. figure 5). This variation in early growth and life history may be explained through differing growth efficiency due to differential metabolic activity (Reid *et al.* 2012; Sloat & Reeves 2014) or through behavioural differences between populations e.g. in feeding aggressiveness (Armstrong et al. 2003; Amundsen & Gabler 2008) or by within-river migration to nursery brooks for better growth opportunities (e.g. Erkinaro & Niemelä 1995). Temporal or microspatial variation in the environment, food availability and predation may maintain growth variation among populations (Amundsen & Gabler 2008; Ward *et al.* 2011; Reid *et al.* 2012; Jonsson & Jonsson 2011). On the other hand, *P*_ST_-*F*_ST_ analysis indicated that, in general, divergence between freshwater traits (other than third year fresh water growth, *Growth*_*FW3*_; Figure 6) generally did not deviate from neutral expectations and therefore variation between the sub-populations may be explained by neutral processes alone. Finally, despite there being significant variation in freshwater growth among populations, there was no difference in freshwater age structures (see Table 3). Several factors may affect freshwater growth and freshwater age similarly (Jonsson & Jonsson 2001), but the lack of observed relationship in this case does not support a mechanistic link between factors resulting in freshwater growth variation among sub-populations, and freshwater age.

It is also of interest to determine if freshwater growth properties may be mechanistically linked to sea age at maturity variation between sub-populations. Larger juvenile size in salmonids is associated with lower mortality (e.g. O’Connell & Ash 1993; Hutchings & Jones 1998; Grover 2005; Jonsson & Jonsson 2011). Therefore, higher freshwater growth of Sub-population 2 individuals may imply lower mortality both in fresh water and during the early marine phase, which predicts a younger age at maturity in Sub-population 2 compared to Sub-population 1 (e.g. Hutchings & Jones 1998; Schaffer 2003). A genetic basis for freshwater growth variation may result in differential optimum age structures in these sub-populations (e.g. Garant et al. 2003), and differences in migratory behaviour may further reinforce post-zygotic isolation between them and help to maintain diversity and population structure within the mainstem. Neither genetic by environment interactions, nor the mechanistic basis of sea age variation is clearly understood in salmonids and therefore resolving this issue awaits further research.

### Implications for conservation

Age at maturity is one of the key traits for the management of Atlantic salmon, as larger multi-sea winter fish are favoured in fisheries. In addition, older age at maturity within a population is correlated with higher genetic diversity and is therefore important for genetic stability of populations and maintaining ecosystem services (Vähä *et al.* 2007; Schindler *et al.* 2010). However, sea age structure is shifting towards younger age classes in many populations (Hansen & Quinn 1998; Niemelä *et al.* 2006; Friedland *et al.* 2009; Chaput 2012; Otero *et al.* 2012). The importance for conservation and management of preserving variation in sea-age within the Teno system has already been recognised (Vähä et al. 2007; Johnston et al. 2014). The results reported here build upon this by providing additional support for targeted preservation programmes, as well as the details necessary for their implementation. Although sea-age has been an obvious target, our assessment of additional phenotypic traits indicated that the phenotypic divergence between the two sub-populations extends beyond sea-age composition, with several growth parameters, including both freshwater and marine growth, differing significantly between sub-populations (Figure 5). Therefore, actions to preserve sea-age variation and/or both sub-populations will serve to preserve diversity in life-history variation expressed during the marine and freshwater phases of the Atlantic salmon life cycle. Detailed population genetic analyses provide further information, by which targeting for sub-population specific preservation is feasible; for example, even though the two sub-populations occur sympatrically throughout the mainstem, Sub-population 2 is more common in the upper reaches. Assessment of historical phenotypic proportions of the sub-populations, which is feasible via the long-term scale archive (Niemelä *et al.* 2006), may be warranted to determine if anthropogenic factors may have altered their life-history make-up and/or sub-population distribution over recent decades and if so, which potential solutions should be proposed.

More generally, our results further indicate that low but significant differentiation revealed by molecular markers can indeed be biologically meaningful, and such subtle, fine scale population differentiation may be overlooked without an integrated analysis of demographic, phenotypic and genetic data. As few within-river genetic studies on salmonids have been conducted with as many genetic markers as used here, it remains to be seen whether Teno River Atlantic salmon represent an exception for the occurrence of such fine scale differentiation in sympatry or whether these findings may be generalized to other large salmon river systems or even more broadly. Likewise, the system appears to be an excellent wild model to study the evolution of life history trade-offs and to improve our understanding of the dynamics of life history evolution both at population and meta-population levels.

## Acknowledgements

We acknowledge the fishermen and women on the Teno River who contributed scales and phenotypic information to the Natural Resources Institute Finland. Scale analyses were carried out by Jari Haantie and Jorma Kuusela. The samples were prepared for SNP genotyping by Katja Salminen and SNP genotyping was conducted at the Centre for Integrative Genomics with the assistance of Matthew Kent and Sigbjørn Lien. Financial support was provided by the Academy of Finland (grants 272836 and 284941).

### Data accessibility

Sampling locations, phenotype data, Structure paramfiles and raw results, and SNP genotypes are available in Dryad doi:10.5061/dryad.7t4n0.

### Author Contributions

J.E., E.N. and P.O. co-ordinated the collection of samples. C.R.P., T.A., J.E., P.O. and S.E.J. designed the study. T.A. analysed the data. T.A. and C.R.P. wrote the first version of the paper. All authors contributed significantly to revisions.

